# Effects of cardiac function alterations on the risk of postoperative thrombotic complications in patients receiving endovascular aortic repair

**DOI:** 10.1101/2022.11.24.517620

**Authors:** Xiaoning Sun, Siting Li, Yuan He, Yuxi Liu, Tianxiang Ma, Rong Zeng, Zhili Liu, Yu Chen, Yuehong Zheng, Xiao Liu

## Abstract

Chronic heart disease (CHD) is a common comorbidity of patients receiving endovascular aneurysm repair (EVAR) for abdominal aortic aneurysms (AAA). The ventricular systolic function determines the hemodynamic environments in aorta, and thus regulating the formation of postoperative thrombus. However, the explicit relationship between ventricular systolic function and EVAR complication of thrombotic events is unknown. Here, we proposed a three-dimensional numerical model coupled with the lumped-elements heart model, which is capable of simulating thrombus formation in diverse systolic functions. The computational results demonstrate that thrombus tended to form on the interior side of the aorta arch and iliac branches, which is consistent with the four patients’ post-operative imaging follow-up. In addition, we found that the thrombus formation has negative correlations with the maximum ventricular contractile force (r=−0.2814±0.1012) and positive correlations with the minimum ventricular contractile force (r=0.238±0.074), whereas the effect of heart rate (r=−0.0148±0.1211) on thrombus formation is not significant. In conclusion, changes in ventricular systolic function may alter the risk of thrombotic events after EVAR repair, which could provide insight into the selection of adjuvant therapy strategies for AAA patients with CHD.

## Introduction

Abdominal aortic aneurysm (AAA) is one of the most common degenerative aortic diseases and is often complicated with various levels of cardiac dysfunction and cardiovascular comorbidities (1). For advanced and/or rapid-growing aneurysms, prompt surgical or endovascular interventions are indicated, among which endovascular aortic aneurysm repair (EVAR) is widely accepted as the first-line treatment option for anatomically feasible cases (2). Cardiac comorbidities may increase the risk of perioperative surgical complications and affect mid-to-long-term prognosis (3). Levels of cardiac function and functional reserve determine the quality of the patients’ tolerance to surgical interventions and the efficacy of postoperative rehabilitation.

Intraluminal thrombus formation is commonly observed not only in major arteries with atherosclerotic and aneurysmal lesions but also, to a lesser degree, intra-prosthetically after endovascular stenting. Despite its lower rate of occurrence, intra-prosthetic thrombus formation may lead to post-operative distal embolism, branch occlusion, and critical end-organ malperfusion (4–6). The formation of intraluminal thrombus and thrombogenic atherosclerotic plaques are believed to be driven by hemodynamic factors. Pathogenic blood flow that generates oscillatory and non-physiological wall shear stress may trigger the accumulation of lipid and pro-thrombotic factors (7). Furthermore, postoperative hemodynamic alterations in the aorta and its major branches are largely determined by the extent of cardiac response and anatomic modifications (8). Individual cardiac functions vary among patients of different ages and physical conditions. However, quantitative data of cardiac functions (cardiac output, stroke volume, etc.) is often unavailable because the measuring process may not be regularly included in the standard protocol of preoperative clinical assessment in real-world scenarios. We noted that only limited and contradicted evidence exists on whether peri- and postoperative use of cardiac supportive drugs such as β-blockers could be beneficial for AAA patients after EVAR in light of aneurysmal sac shrinking and clinical outcomes (9–11). The effects of variations in cardiac function on the risk of intraluminal and intra-prosthetic thrombus formation are yet to be further elaborated.

By modeling the process of hemodynamics and mass transport in the coagulation cascade, the patient-specific patterns of formation and distribution of intraluminal/intra-prosthetic thrombus formation can be simulated or predicted with acceptable accuracy (12, 13). Based on different thrombus initialization mechanisms, numerous computational models for thrombus formation have been developed in literature (14–20). However, the explicit effects of cardiac functions on thrombus formation were not yet well addressed. In order to simulate the coupling between the cardiac function and the hemodynamics environments, we used the lumped parameters-3D coupled models(21, 22). These models are capable of reflecting the heart function and its effect on the blood flow environment in lumped parameters model with acceptable accuracy in a significantly shorter time. We can use the pressure and velocity profile output from the 0D heart model as inlet conditions for 3D continuum thrombus model. In this pilot study, by coupling a lumped parameters model of cardiac output and aortic/peripheral outflow resistance with a 3D numerical mass transport model of thrombus formation, we focused on investigating the relation of cardiac functions and the predicted risk of thrombus formation in the aorta and/or endograft of 4 patients who underwent EVAR. Relative risks for thrombus formation were identified using machine-learning algorithms.

## Methods

### Imaging acquiring and 3D model reconstruction

Four patients diagnosed with AAA who received EVAR and regular follow-ups at the department of vascular surgery of Peking Union Medical College Hospital were included. DICOM format of CTA images of each patient from preoperative and at least 6 months post-operative was obtained. Mimics 12.0 (Materialise, Belgium) and CRIMSON (25) were used to reconstruct 3D models of the aorta from ascending aorta to aortoiliac bifurcations, with preservation of major branches in the arches and the thoracoabdominal segments (Figure 1a). Models were subsequently smoothed with WRAP (v2017, Geomagic) and loaded into SOLIDWORKS (v18, Solidworks Co.). Vessel outlets were trimmed and extended sufficiently to allow for the fully developed flow pattern. The geometric model was meshed with a tetrahedral mesh using ICEM (v17.0, ANSYS, Inc.). The global meshing size was chosen at 0.02 mm to accommodate the smallest components. Elements of the boundary layers were set to be hexahedral with a primary thickness of 0.001 mm and would gradually develop over 5 layers (Figure 1b). Under steady-state conditions, grid independence is considered to be achieved when the mean difference between WSS and platelet concentrations in two continuous simulations is less than 1%. The study followed the Helsinki Declaration and was approved by the ethical committee of Peking Union Medical College Hospital. Written informed consent was obtained from all participants.

**Figure 1.**
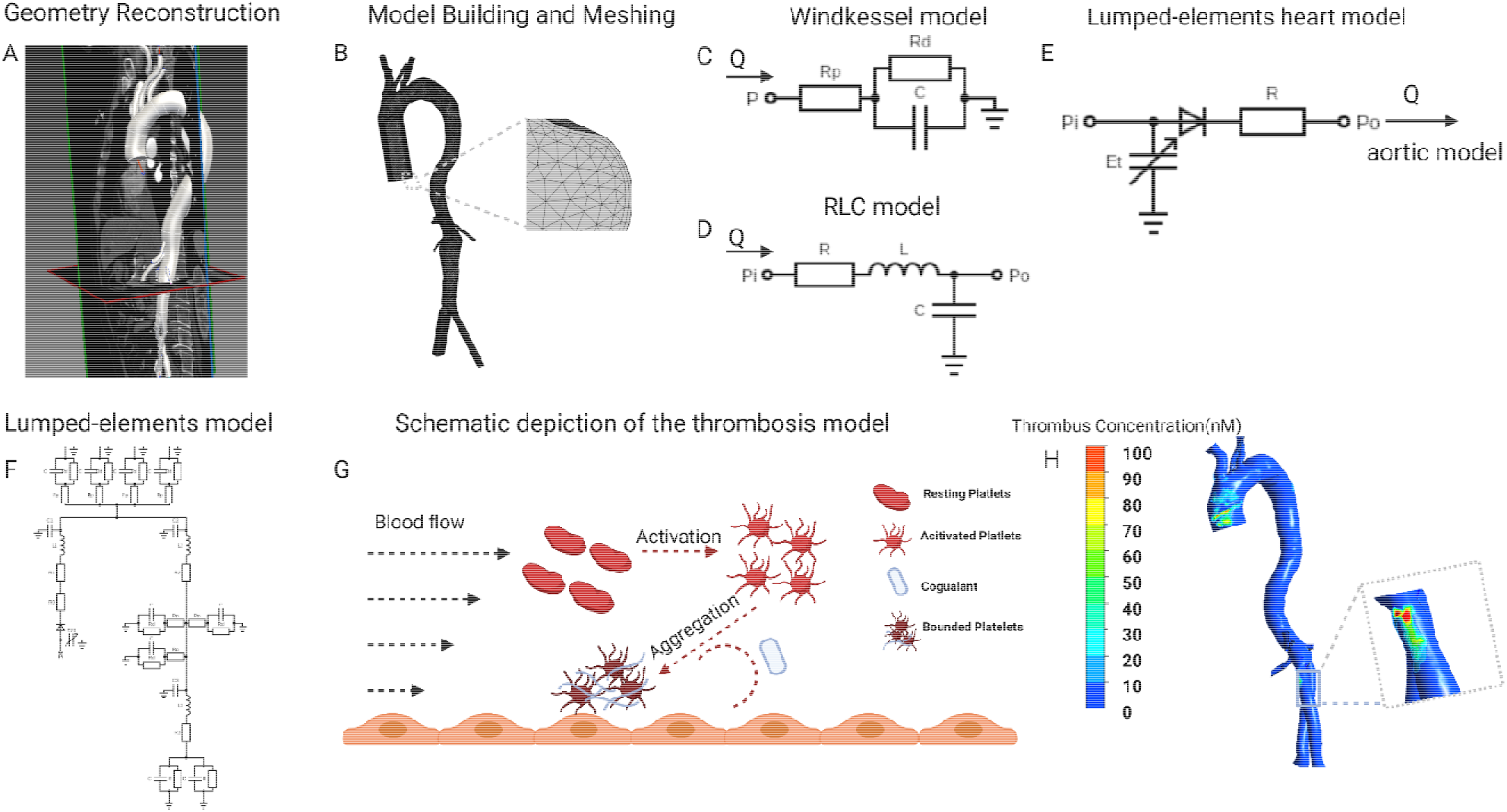
Flowchart of patient-specific modeling. **A**, Geometry of postoperative aorta was modeled from CTA data, starting from the ascending aorta to bilateral common iliac arteries. **B**, Meshing the model and vessel outlets were sufficiently extended to allow fully developed flow boundary. **C, D**, Building the Windkessel model, lumped-param ter aortic model based on the clinical measured blood flow pressure and velocity. **E**, The lumped-elements model representing the heart was coupled to the entrance node in the lumped-elements aortic model. **F**, Coupling the three kinds of lumped-parameter model to give out the aortic blood flow velocity in different heart functions. **G**, The simulation of thrombosis formation involving the activation of resting platelets (RPs), the deposition of coagulant (C) and the aggregation of activated platelets (APs) to form the bounded platelets (BPs). **H**, Thrombus concentration profile of one patient. CTA, computed tomography angiography. The figure was created with BioRender.com

### Mathematical modeling for 0D model

The aorta was separated into three sections and represented by three RLC elements. The outlets of the aorta are modeled as the three elements Windkessel model(26). The parameters in the RLC element and Windkessel model were calculated by solving the differential equations based on the blood flow pressure and velocity. In addition, a function imitating ventricular systole is used in the lumped parameter model of the heart module. Through animal tests, Suga and Sagawa(27) developed a ventricular pressure-volume relationship that can be expressed as a time-varying function E(t), as indicated in Equation (1):

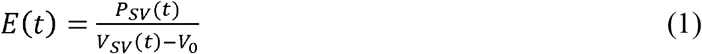

where E(t) is the time-varying function (mmHg/ml), P_sv_(t) is the time function of ventricular pressure (mmHg), V_sv_(t) is a time function of ventricular volume (ml), and V_0_ is the ventricular reference volume (ml), which is the theoretical volume relative to ventricular zero pressure. Boston(28) proposed a mathematical fit to determine the function of ventricular systole according to equation (2):

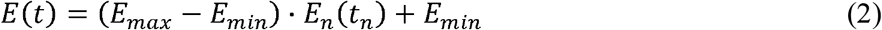

where E_max_ refers to the ventricular pressure-volume relation at end-systole, E_min_ refers to the ventricular pressure-volume relation at end-diastolic. E_n_(t_n_) represents the normalized function of ventricular elasticity, which is described as the Hill equation as follows(29):

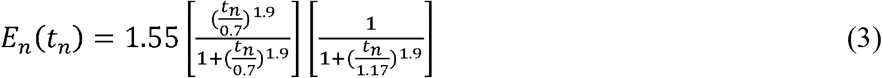

where t_n_ is t/T_max_, and the T_max_ can be calculated from the cardiac cycle:

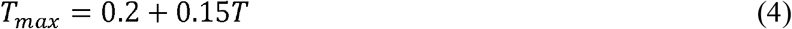

### Mathematical modeling for 3D simulation of thrombus formation

In this article, we simulate the 3D formation of thrombus by solving the convection-diffusion-reaction equations. The thrombus formation is tracked through the formation of bounded platelets (BPs), which was induced by localized high concentration of activated platelets (APs), coagulation enzymes (C), and prolonged flow residence time (RRT). The schematic of the thrombus formation is illustrated in Figure 1 g. We only show the general form of the equation here, and the complete reactions and parameters adapted from the published models (13, 30, 31) are explained in detail in Table S2 and Table S3:

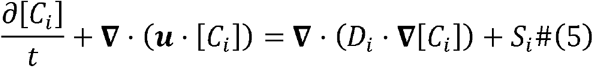

where [*C_i_*], is the concentration of species *i; u* represents the blood flow velocity vector; *D_i_* refers to the diffusivity of species i in blood; and *S_i_* is a local reaction source term for species i. The species considered in this model include rested platelets [RPs], activated platelets [APs], coagulation enzymes[C], and flow residence time [RT].

The bounded platelets (BPs) attached to the reactive surface have neither convective nor diffusive motion; therefore, in this case, Equation (6) is simplified and the deposition of platelets are governed by the following concentration rate equations:

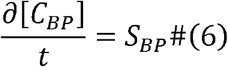

where *S_BP_* is the local generation rate of bounded platelets (BPs).

Considering the resistance of bounded platelets to blood flow, the momentum source term was added to the Navier-Stokes equations, and the viscosity coefficient was modified. Equation (7) shows the modified Navier-Stokes equations. and Eq (8) and (9) show the modified viscosity and source term:

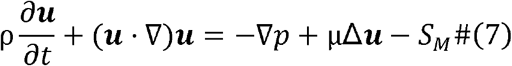

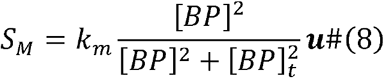

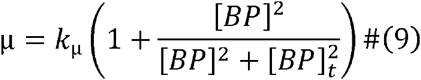

where ρ represents the blood density equals 1050*kg/m^3^*, *u* represents the blood flow velocity vector, *p* represents blood pressure, μ represents the blood viscosity, and *S_M_* represents the local source term of flow momentum. Detailed value of coefficient *k_m_*, *k_μ_* can be found in supplement material (Supplementary Table S3).

### Multiscale simulation methods and procedures

Our model coupled the 3D simulation with the lumped-elements model. First, based on the clinically measured blood flow pressure and velocity(13), we transformed the aortic model from 3D to lumped elements and coupled the boundary nodes to Windkessel elements(32) (Figure 1c, d, detailed parameters in Supplementary Table S1). Subsequently, the lumped-elements model representing the heart was coupled to the entrance node in the lumped-elements aortic model (Figure 1e). We first adjusted the heart model’s parameters so that the averaged cardiac output and aortic inlet blood pressure levels matched the clinically determined values. To mimic the various heart functions, we swept the parameters E_max_ (end-systolic elastance), E_min_ (end-diastolic elastance), and T (heart rate) from 10% to −10% compared to the reference values and got the corresponding aortic blood flow velocity in different heart functions. Finally, we simulated the formation of thrombus in the 3D aorta under various cardiac functions, the boundary conditions of which are derived from the blood flow velocity at the boundary nodes in the lumped-elements model (Figure 1f, g). The 3D simulations of blood flow and thrombus formation were performed in FLUENT (v17.0, ANSYS, Inc.) with user-defined functions, whereas the lumped-elements simulations were performed in MATLAB (R2021b, MathWorks, Inc.) The results were analyzed and visualized using CFD-Post (v17.0, ANSYS, Inc.).

### Statistical Analysis

Data were analyzed using Prism software (v9.0, GraphPad Prism, Inc). Statistical significance for samples was determined using two-tailed unpaired Student’s t-test. Data were considered statistically significant if P < 0.05. The Parallel coordinate analysis and the Multiple correspondence analysis are performed in MATLAB (R2021b, MathWorks, Inc.).

## Results

### Demographic characteristics and clinical verification of thrombus formation region

The baseline information of 4 patients were shown in table 1. There were 3 male and 1 female patients, with an average age of 74.3 ± 7.6 years. Pre-operatively, laboratory results indicated normal cardiac and renal function for the 4 subjects. All 4 patients underwent the operation successfully, and follow-up CTA suggested patency of stents and visceral branches. Concerning thrombus formation, CTA results in Figure 2 (A1-D1) illustratrd that thrombus was found at the root of the left subclavian artery for patient number 3. For patient number 1, 2, and 4, CTA images all showed thrombus within the iliac stents. Thrombus was not observed at other parts of the arterial or stent model from clinical images. These regions with thrombus formation were defined as the region of interests (ROI) in the following analysis.

**Table 1.** Clinical information and total BP levels of patients under different heart conditions.

|  | Patient 1 | Patient 2 | Patient 3 | Patient 4 | Average |
| --- | --- | --- | --- | --- | --- |
| Age, years | 80 | 70 | 64 | 83 | $74.3 \pm 7.6$ |
| Sex | Male | Female | Male | Male | - |
| Hypertension | Yes | Yes | Yes | Yes | - |
| Diabetes | No | Yes | No | No |  |
| Smoking | Yes | No | Yes | No | - |
| Laboratory results |  |  |  |  |  |
| LVEF% | 68 | 74 | 67 | 77 | $71.5 \pm 4.2$ |
| NT-proBNP, pg/ml | 104 | 112 | 81 | 77 | $93.5 \pm 14.8$ |
| D-dimer, mg/L | 6.31 | 5.3 | 1.67 | 1.41 | $3.7 \pm 2.2$ |
| HGB, g/L | 123 | 121 | 150 | 120 | $128.5 \pm 12.5$ |
| Creatinine, $\mu\text{mol/L}$ | 78 | 70 | 93 | 111 | $88.0 \pm 15.6$ |
| Follow-up, months | 35.5 | 47.7 | 34.8 | 11.9 | $32.5 \pm 14.9$ |

BP, bound platelets; $E_{\text{max}}$ , maximum elastase; $E_{\text{min}}$ , minimum elastase; LVEF, left ventricular ejection fraction;
NT-proBNP, N-terminal pro-brain natriuretic peptide; HGB, hemoglobin.

**Figure 2.**
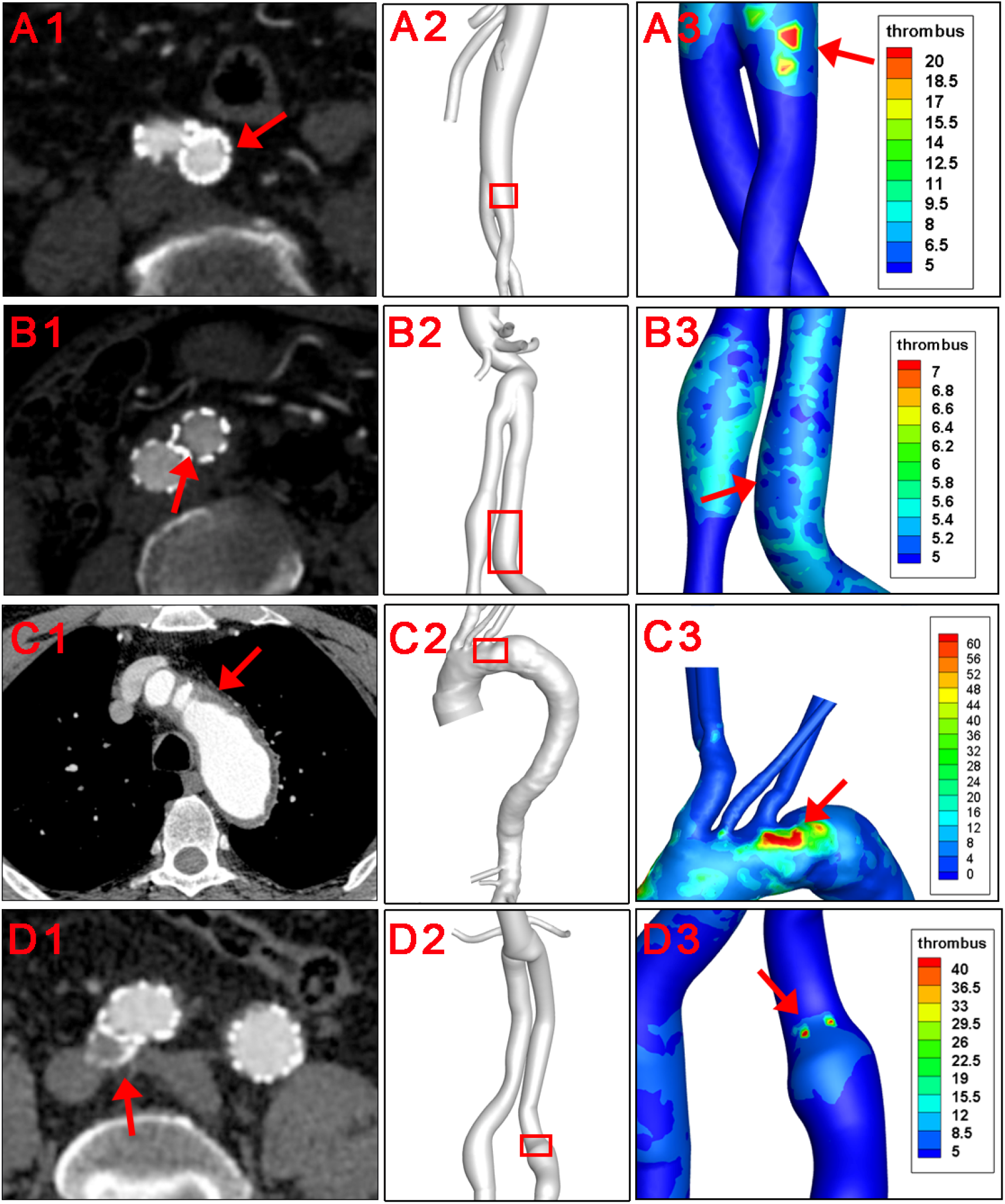
Clinical verification of the thrombus forming model. Red arrows and red squares indicated places of thrombus formation on CTA images and corresponding 3D models.

The hemodynamic parameters and BP levels at ROI were demonstrated in Figure 2 and Figure 3. The TAWSS, oscillatory shear index (OSI), and relative residence time (RRT) profiles of 4 models were illustrated in Figure 3. In general, TAWSS was highest at the visceral branch level in all 4 patients, which indicated higher frictional force exerted on these vessel areas by the blood flow and prevented activated platelets from binding to coagulant to form thrombus in our model. Descending aorta and iliac branches had higher RRT, which indicated a higher degree of retention for a substance in the bloodstream at the region and induced the development of thrombus in our model (See supplementary material). RRT also increased at the oversizing area for the iliac branches. However, zooming in on the ROI in Figure 2 did not indicate distinct abnormal local CFD parameters. In addition, no significant difference in TAWSS, OSI, and RRT was observed for the ROI compared to the average levels of the whole model (right channel in Figure 3). On the other hand, patterns of BP distribution from the thrombus model simulation (Figure A2-D2) corresponded well with the thrombus results from ROI.

**Figure 3.**
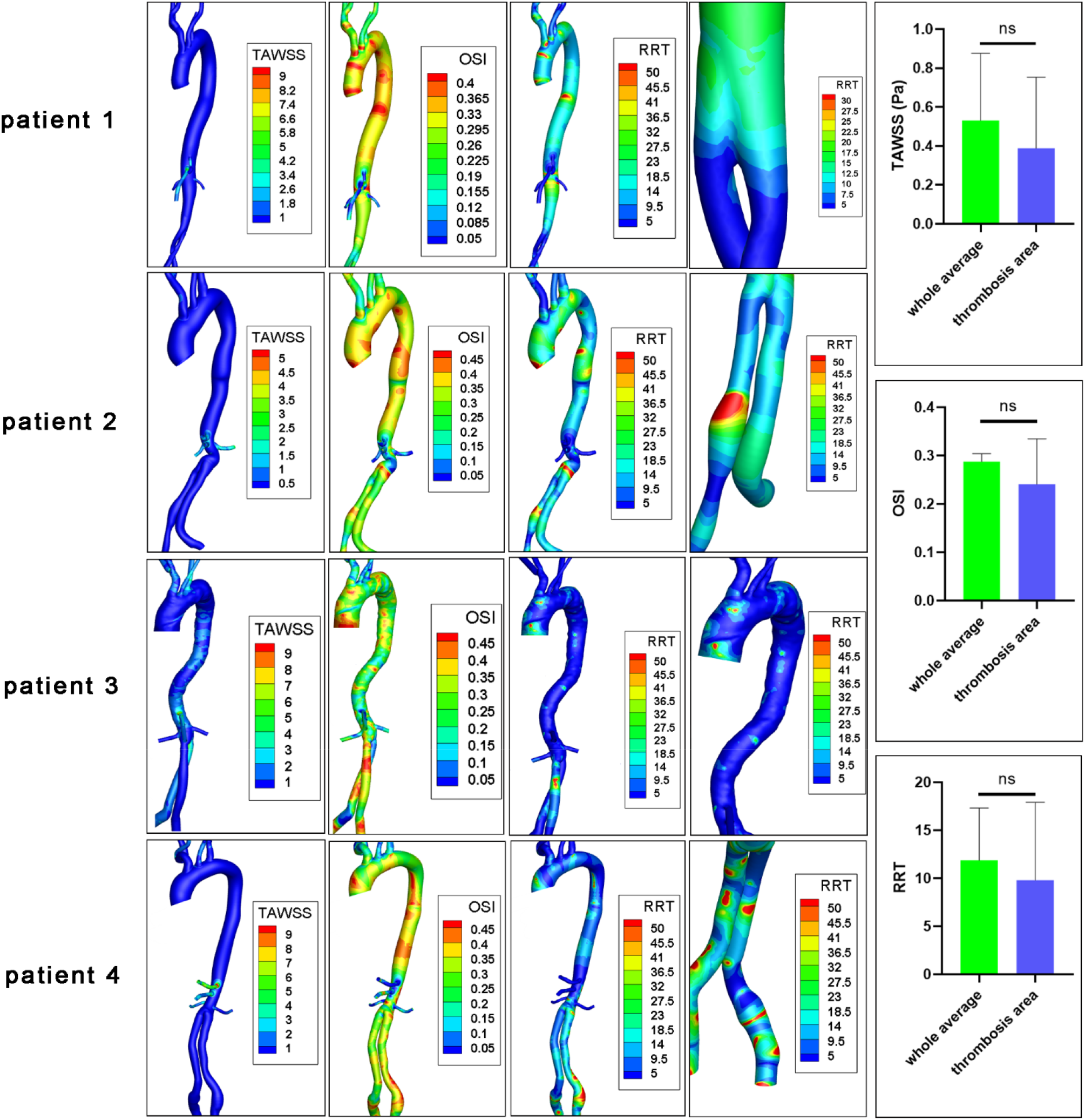
CFD characteristics of patients’ original models. TAWSS, time average wall shear stress (Pa); OSI, oscillatory shear index; RRT, relative residence time (Pa^−1^); ns, non-significant. The fourth columns indicated RRT levels at ROIs.

### Impact of cardiac function alterations on thrombosis

The impact of heart function alteration on thrombus formation in the ROI as well as the aortic inlet velocity were shown in Figure 4. For patient 1, 2, and 4, the iliac branches were shown, while the arch and descending aorta was illustrated for patient 3.

**Figure 4.**
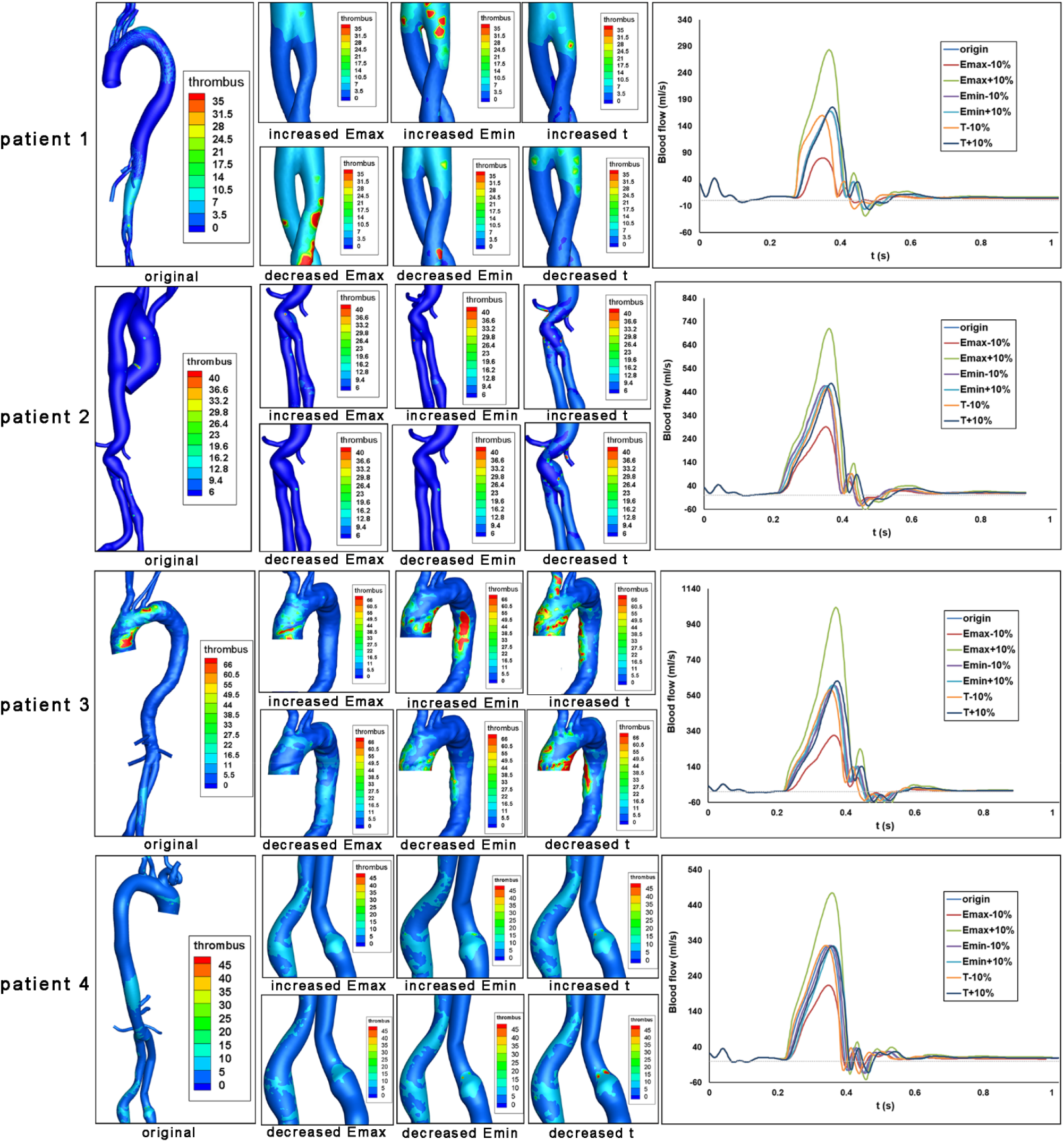
Impact of heart function alteration on thrombosis at critical positions. Bounded platelets (BP) levels under different heart conditions were shown for each patient. Aortic blood flow values over a cardiac cycle were also shown.

The aortic velocity profile demonstrated that the magnitude and shape of the blood flow profile are affected by the heart function parameters of E_max_, E_min_, and T. Then we analyzed the formation of thrombus, which was induced by the altered heart functions. First of all, the increased E_max_ (end systolic elastance, representing maximum ventricular contractile force) could maintain or alleviate thrombus formation levels in the ROI of four patients. However, the effects of decreased E_max_ are heterogeneous. For patient 1, a decrease in E_max_ would result in an increase in thrombus development at the iliac branches. However, for patients 2-4, the lower Emax would result in less thrombus development at the arch and descending aorta. In addition, the increased E_min_, which represents ventricular compliance, would maintain or exacerbate the thrombus formation in the ROI of four patients. The decreased E_min_ would maintain or alleviate the thrombus formation in these regions. Furthermore, the influences of heart rate (T) on thrombus formation are less significant.

We evaluated the area-averaged BP concentration and area-averaged TAWSS (traditional indicator) of the four patients to quantitatively highlight the associations between heart function and thrombus formation in ROI. We discovered that TAWSS is largely affected by Emax level, while Emin and T have little impact by showing the Parallel coordinate charts classified by the heart function (Emax, Emin, T ranges from +10% to −10% compared with the reference condition). In addition, we found that increasing Emax and decreasing Emin could reduce overall thrombus formation levels (Figure 5a–5b), which is in agreement with the qualitative analysis. Besides, both the increase and decrease of heart rate (T) would exacerbate the thrombus formation level (Figure 5c). The effects of Emin and t on TAWSS, however, were not significant, indicating the limitations of using hemodynamic features to predict thrombus development. Then, we performed the multidimensional analysis method to quantitatively reveal the correlations among E_max_, E_min_, T, and thrombus formation. We found that the BP concentration was negatively correlated with E_max_ (mean±SD: r=−0.2814±0.1012, represents the average correlation coefficient at different segments of the aorta for four patients), and was positively related with E_min_ (r=0.238±0.074) (Figure 5c). However, there was a lack of correlation between heart rate (T) and BP concentration (r=− 0.0148±0.1211). In addition, we found that the ROI (iliac in patient 1,2,4 or arch in patient 3) showed the highest correlations with Emax and Emin, which indicated that the thrombus formation in these regions was mostly sensitive to the alteration in heart function.Collectively, these data indicated that the increase in E_max_ and decrease in E_min_ would both inhibit thrombus formation, and the effect of T was not significant.

**Figure 5.**
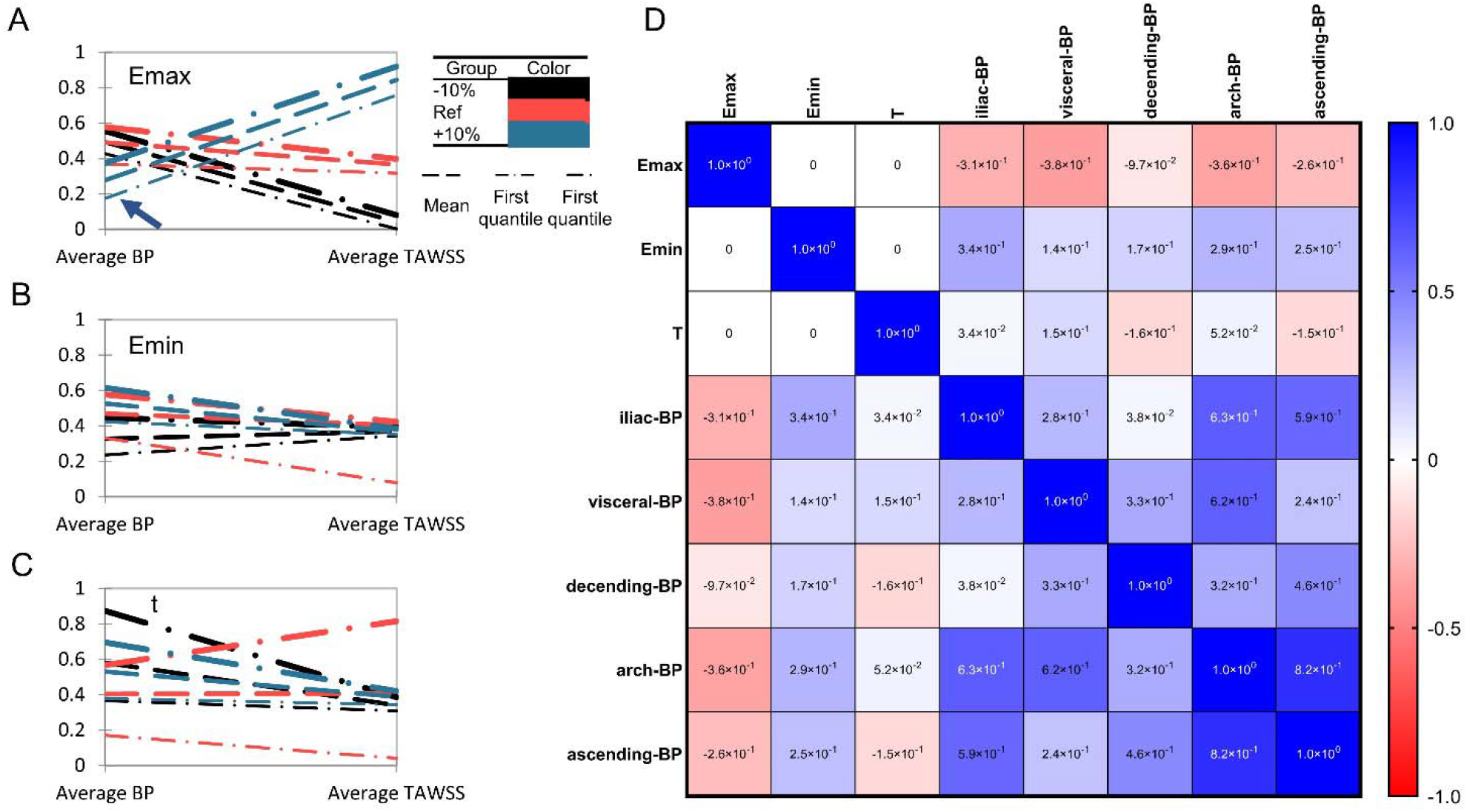
Heart function alteration and change of thrombosis or TAWSS at different aortic parts. A-C. Parallel coordinate charts of BP concentration and TAWSS classified by heart function (E_max_, E_min_, T ranges from +10% to −10% compared with the reference state). BP concentration and TAWSS (Pa) have been rescaled ranges from 0 to 1. The arrowhead in Figure 5A indicates the significant role of E_max_ on thrombus formation. D. Multiple correspondence analysis (MCA) of E_max_, E_min_, T, and BP concentration in different aortic sections.

## Discussion

Endograft mural thrombus accumulation had reported rates up to 33% and was detected as early as 1 week after EVAR (4). It might be due to cytokine and prothrombotic factors released from the intramural thrombus of the aneurysm triggered by the operation. Whether the risk of long-term thrombotic events could be increased is debatable (33). Mestres et al. reported that endograft mural thrombotic deposits were related to device occlusion during 24 months of follow-up (34), while Melson et al. did not find a significant association (35). Previous studies on CFD-based thrombus simulation mainly focused on the intra-luminal thrombosis in the sac of AAA. Abnormal wall shear stress, platelet activation, vortical structures, and morphological parameters were all found to play a potential role (36–39). Regarding intra-prothetic thrombosis, Nauta and colleagues investigated the alteration of PLAP in one patient receiving 3 virtual interventions including TEVAR (40). Liu et al. explored the TAWSS, OSI, and RRT levels in 3 patients who underwent multibranched endovascular repair (41). Most of these studies referred to previous works or idealized values for boundary blood flow and pressure profiles.

Thrombosis involves complex interlinked interactions between platelets, coagulation cascades, and the vascular wall (42). In this study, traditional hemodynamics parameters including WSS, OSI, and RRT did not precisely identified regions of thrombus formation, probably due to the lack of reflection of this complex reactions. In a former study, we used a numeric thrombus prediction model to calculate the vascular remodeling and thrombotic events in one patient receiving hybrid repair for the middle aortic syndrome, which was consistent with follow-up images (13). The continuum-macroscopic scale model could capture the clotting patterns taking both activated platelets, local hemodynamic conditons, and residence time into account (12). Herein, this numeric model was applied to 4 patients receiving EVAR. In line with CTA results, thrombus formation was observed at iliac branches in 3 patients and at the opening of LSCA in 1 patient. Iliac endografts are commonly reported places for thrombosis, and the aorto-uni-iliac configuration was confirmed as a risk factor for intra-prosthetic thrombus (IPT) deposit in a meta-analysis (5). The low-density mural thrombus in the descending aorta, on the other hand, may originate from the disruption of vulnerable atherosclerotic plaques (43). Our result suggested that this numeric model was suitable for both intra-luminal and intra-prosthetic thrombosis prediction.

Previous works investigating the relationship between cardiac function and EVAR complications were mainly retrospective cohorts or reviews (10, 44). To the best of our knowledge, this is the first attempt at using a cardiac numeric model for post-EVAR patients. Lumped heart models used simple parameters such as resistance and capacitance to simulate the relaxation, filling, contraction, and ejection phases of the heart, and have been widely studied for their interaction with the arterial vessel systems (45). In this study, we used a 0-D cardiac model adapted from Kim et al. to simulate the change in heart function especially ventricular pressure/volume depending on the time-varying elastance curve (21). Emax represented the maximum ventricular contractility, Emin represented ventricular compliance or end-diastolic pressure, and t stood for the cardiac cycle. Heart failure with reduced ejection fraction (HFrEF) or with preserved ejection fraction (HFpEF) could be mimicked under the circumstances. The influence of cardiac parameters on thrombosis was reflected by altered BP levels.

Concerning the actual thrombus forming area, both elevated or reduced E_min_ and t promoted thrombus formation, yet changes of Emax induced distinct results. It should be noted that BP levels in patients 1 and 3 were higher and more concentrated around the actual thrombus-forming area. Despite thrombus deposition in the local iliac stent, the BP in patients 2 and 4 had relatively low absolute value and occupied a more extensive spatial range. Morphological factors such as increasing diameter because of stent oversize may have a stronger influence on thrombosis in these patients, and the impact of cardiac parameter alteration should be cautiously interpreted. The heterogeneous results from different patients emphasized the importance of individualized analysis.

Quantitation calculation of BP for general aortic parts revealed that Emax was negatively associated with thrombus formation while E_min_ was positively related. It has been reported that male AAA patients had reduced ventricular systolic and diastolic function (46). Our result indicated that for patients with CHD (both HFrEF and HFpEF) in clinical practice, we should pay more attention to their increased risk of thrombus formation after EVAR. Additionally, preservation of maximum ventricular contractile force and stabilization of ventricular compliance or heart rate was also a crucial consideration when selecting supporting drugs.

This study has some limitations. Patient-specific outflow data of each aortic branch may not always be available, especially for supra-arch branches in AAA patients, and assumptions were made for blood distribution in a limited number of branches according to the literature. The simulation model for thrombus formation could estimate the tendency but not the speed of thrombosis in the vessel model. The simulation process should be applied in a larger patient cohort with multiple follow-up images to validate our findings. Besides, we ignored the arterial deformation in the 3D calculation. Future studies may combine fluid-structure interaction (FSI) models with mass transport models of thrombus to more realistically predict thrombus formation in deformed vessels(47, 48).

## Conclusion

In conclusion, changes in ventricular contractile force may alter the risk of thrombus formation for post-EVAR patients, whereas the effect of heart rate is not significant. Postoperative cardiac support recommends drugs that support maximum ventricular contractile force, maintain left ventricular pressure, and stabilize heart rate.

## Author Contributions

X.S, S.T., Y.H., Y.L., and T.M. led the modelling and calculation of thrombus formation. X.S., T.M. Y.Z., X.L. conceived, designed, and led the interpretation of the project. X.S, S.T., R.Z., Z.L. provides and analyses the clinical data. Y.C. constructed the lumped-elements model. All authors contributed to the writing of the manuscript.

## Acknowledgments

None.

