## supplement material for "Effects of cardiac function alterations on the risk of postoperative thrombotic complications in patients receiving endovascular aortic repair"

Supplementary Material

### 1 Supplementary Data

**Boundary condition acquisition**

EKG-gated Doppler ultrasound (D-US) was applied to measure mean blood flow velocity in both subclavian artery (SCA), both common carotid arteries (CCA), celiac axis (CA), inferior mesenteric artery (IMA), both renal arteries, and both common iliac arteries. Site for Doppler sonogram acquisition was several centimeters distal to the origin of each branch vessel, avoiding any abrupt torsion or tapering. Transient mean flow velocity of vessel branches was calculated by intrinsic software of the ultrasound scanner (iUElite, Philips; Netherlands), and was modified and smoothed using MATLAB (vR2018a, MathWorks; Natick, USA). Representative flow velocity was obtained between two adjacent R-wave as marked in the EKG leads, thus flow velocities of outlet vessels were synchronized accordingly. D-US derived transient velocity profile was assigned to each outlet branch. For two patients who did not have adequate D-US profiles of each vessel branch, assumptions were made for blood distribution according to the literature (1). Pressure profile at the aortic inlet was acquired from intra-operative Invasive arterial pressure monitoring. The parameters in the RLC element and Windkessel model were calculated by solving the differential equations based on the blood flow pressure and velocity.

### 2 Supplementary Figures and Tables

TABLE 1 Partial differential equations of lumped-elements

| **Partial differential equations of lumped-elements** | | | | | | | | |
| --- | --- | --- | --- | --- | --- | --- | --- | --- |
| **Lumped-elements model** | | | **The position used for the model** | | **Schematic** | | | **Equations** |
| The three-elements Windkessel model | | | ca/lcca/lra/lsca/rcca/rra/rsca/sma | | 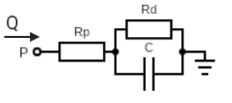 | | | $\frac{dP}{dt}+\frac{P}{R_{d}C}=\frac{Q}{C}(1+\frac{R_{p}}{R_{d}})+R_{p}\frac{dQ}{dt}$ |
| The two-elements Windkessel model | | | lcia/rcia | | 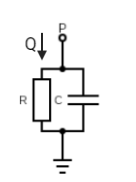 | | | $\frac{dP}{dt}+\frac{1}{RC}P=\frac{Q}{C}$ |
| RLC model | | | Each section of aorta | | 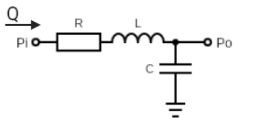 | | | $P_{i}-P_{o}=R\cdot Q+L\frac{dQ}{dt}$ |
| **Details in the lumped-elements model** | | | | | | | | |
| **Lumped-elements model** | | | **Patient ID** | | | | | |
|  |  |  | 1 | 2 | | 3 | 4 | |
| Windkessel model | Rp(pa·s/m^3^) | ca | 2.37E+08 | 2.07E+08 | | 1.00E+08 | 1.02E+08 | |
|  |  | lcca | 2.16E+08 | 3.5E+08 | | 2.01E+08 | 1.96E+08 | |
|  |  | lcia | 2.38E+08 | 2.79E+09 | | 9.44E+08 | 7.89E+09 | |
|  |  | lra | 3.92E+08 | 1.1E+09 | | 1.23E+08 | 6.70E+08 | |
|  |  | lsca | 1.71E+08 | 1.3E+08 | | 3.46E+08 | 4.16E+08 | |
|  |  | rcca | 2.46E+08 | 2.43E+08 | | 3.12E+08 | 2.24E+08 | |
|  |  | rcia | 2.03E+08 | 7.23E+08 | | 6.36E+08 | 6.67E+09 | |
|  |  | rra | 4.57E+08 | 1.2E+09 | | 1.66E+08 | 4.62E+08 | |
|  |  | rsca | 4.36E+08 | 1.97E+08 | | 1.42E+08 | 3.74E+08 | |
|  |  | sma | 6.41E+07 | 3.8E+08 | | 1.15E+08 | 1.73E+08 | |
|  | Rd(pa·s/m^3^) | ca | 2.37E+09 | 2.07E+09 | | 1.00E+09 | 2.07E+10 | |
|  |  | lcca | 2.16E+09 | 3.46E+09 | | 2.01E+09 | 1.96E+09 | |
|  |  | lcia | 2.38E+09 | 2.79E+10 | | 9.44E+09 | 7.89E+10 | |
|  |  | lra | 3.92E+09 | 1.11E+10 | | 1.23E+09 | 1.82E+11 | |
|  |  | lsca | 1.71E+09 | 1.26E+09 | | 3.46E+09 | 2.68E+10 | |
|  |  | rcca | 2.46E+09 | 2.43E+09 | | 3.12E+09 | 2.24E+09 | |
|  |  | rcia | 2.03E+09 | 7.23E+09 | | 6.36E+09 | 6.67E+10 | |
|  |  | rra | 4.57E+09 | 1.23E+10 | | 1.66E+09 | 7.60E+10 | |
|  |  | rsca | 4.36E+09 | 1.97E+09 | | 1.42E+09 | 2.40E+10 | |
|  |  | sma | 6.41E+08 | 3.78E+09 | | 1.15E+09 | 5.29E+10 | |
|  | C(pa·s/m^3^) | ca | 1.12E-10 | 2.63E-10 | | 6.02E-11 | 2.93E-10 | |
|  |  | lcca | 1.16E-10 | 4.07E-10 | | 7.83E-11 | 1.14E-10 | |
|  |  | lcia | 8.43E-10 | 3.69E-09 | | 1.37E-10 | 4.59E-10 | |
|  |  | lra | 6.10E-11 | 6.98E-11 | | 9.22E-11 | 2.96E-11 | |
|  |  | lsca | 5.14E-10 | 2.24E-09 | | 1.89E-10 | 2.22E-10 | |
|  |  | rcca | 1.03E-10 | 5.21E-10 | | 4.80E-11 | 9.97E-11 | |
|  |  | rcia | 9.83E-10 | 6.89E-10 | | 2.12E-10 | 5.21E-10 | |
|  |  | rra | 5.27E-11 | 6.18E-11 | | 6.00E-11 | 5.09E-11 | |
|  |  | rsca | 3.88E-10 | 2.87E-09 | | 4.67E-10 | 2.48E-10 | |
|  |  | sma | 1.27E-10 | 2.52E-10 | | 1.55E-10 | 2.13E-10 | |
| RLC model | R(pa·s/m^3^) | | 2.48E+06; 4.07E+06; 9.31E+06 | 1.79E+06; 8.53E+06; 1.87E+07 | | 2.48E+06 \| \|; 4.07E+06 \| \|; 9.31E+06 \| \| | 6.67E+09  4.07E+06  9.31E+06 | |
|  | L(pa·s^2^/m^3^) | | 8.43E+05; 1.80E+06; 4.47E+06 | 9.74E+05; 3.79E+06; 5.94E+06 | | 8.43E+05 \| \|; 1.80E+06 \| \|; 4.47E+06 \| \| | 8.43E+05  1.80E+06  4.47E+06 | |
|  | C(m^3^/pa) | | 1.38E-09; 1.50E-09; 3.00E-10 | 5.09E-09; 6.64E-09; 2.22E-08 | | 1.38E-09 \| \|; 1.50E-09 \| \|; 3.00E-10 \| \| | 4.59E-10  1.50E-09  3.00E-10 | |

TABLE 2 Species units, diffusive terms, source terms and initial condition

| **Species [C_i_]** | **Di** | **S_i_** | **Initial Condition** | **Unit** |
| --- | --- | --- | --- | --- |
| [RP] | $D_{t}+\alpha\dot{\gamma}$ | -$k_{1}$[AP][RP]- $k_{2}$[C][RRT] | 19 | nM |
| [AP] | $D_{t}+\alpha\dot{\gamma}$ | $k_{1}$[AP][RP]+ $k_{2}$[C][RRT] | 1 | nM |
| [BP] | N/A | $k_{BP}[AP]\frac{\left[ RRT \right]^{2}}{\left[ RRT \right]^{2}+\left[ {RRT}_{t} \right]^{2}}\frac{\left[ C \right]^{2}}{\left[ C \right]^{2}+\left[ C_{t} \right]^{2}}\frac{\overline{{\gamma_{t}}^{2}}}{\overline{\gamma^{2}}+\overline{{\gamma_{t}}^{2}}}$ | 0 | nM |
| [C] | $D_{c}\frac{\overline{{\gamma_{t}}^{2}}}{\overline{\gamma^{2}}+\overline{{\gamma_{t}}^{2}}}$ | $k_{c}\frac{\left[ C \right]^{2}}{\left[ C \right]^{2}+\left[ C_{t} \right]^{2}}\frac{\overline{{\gamma_{t}}^{2}}}{\overline{\gamma^{2}}+\overline{{\gamma_{t}}^{2}}}$ | 0 | nM |
| [RT] | $D_{RT}$ | 1 | 0 | 1 |

$\dot{\gamma}$is the local shear rate.

TABLE 3 Thrombus formation model parameters

| **Parameter** | **Symbol** | **Value/Expression** | **Unit** |
| --- | --- | --- | --- |
| Platelets thermal diffusivity | $D_{t}$ | $1.6\times{10}^{-13}$ **^(2)^** | $m^{2}/s$ |
| Platelets diffusive coefficient | $\alpha$ | $7\times{10}^{-13}$ **^(2)^** | $m^{2}$ |
| Platelets reaction source term coefficient 1 | $k_{1}$ | 0.5 **^(3)^** | $s^{-1}$ |
| Platelets reaction source term coefficient 2 | $k_{2}$ | 0.15 **^(3)^** | $s^{-1}$ |
| Platelets bounded coefficient | $k_{BP}$ | $2\times{10}^{-8}$ | $nM/s$ |
| Relative residence time | RRT | $\frac{\left\vert RT_{t=T}-RT_{t=0} \right\vert}{T}$ | 1 |
| Relative residence time threshold | $RRT_{t}$ | 0.9 | 1 |
| Coagulant diffusivity | $D_{c}$ | ${10}^{-8}$ | $m^{2}/s$ |
| Coagulant kinetic constant | $k_{c}$ | 200 | $nM/s$ |
| Shear rate threshold | $\overline{\gamma_{t}}$ | 50 | $s^{-1}$ |
| Coagulant concentration threshold | $C_{t}$ | 10 | $\mathrm{nM}$ |
| Diffusivity of residence time (RT) | $D_{RT}$ | $1.14\times{10}^{-11}$ | $m^{2}$ |
| Viscosity coefficient | $k_{\mu}$ | 0.0035 | $kg/m^{2}s^{2}$ |
| Moment source term coefficient | $k_{m}$ | ${10}^{7}$ | $kg/m^{3}s$ |

TABLE 4 Boundary condition

| Boundary | APs | RPs | RT | C | BPs |
| --- | --- | --- | --- | --- | --- |
| Inlet | $C_{AP}=1$ | $C_{AP}=19$ | RT = 0 | C = 0 | BP = 0 |
| Outlet | Flux = 0 | Flux = 0 | Flux = 0 | Flux = 0 | Flux = 0 |
| Wall | Flux = 0 | Flux = 0 | Flux = 0 | Flux1 | Flux = 0 |

Flux1 is described as:

$$D_{c}\frac{\partial C}{\partial n}=\left\{ \begin{aligned} 20nM\cdot m/s & if TWASS<0.2Pa and BPs<200nM \\ 0 Otherwise \end{aligned} \right.$$

TABLE 5 Wall shear stress in mesh independency analysis

| Coarse Fine | | | | | | |
| --- | --- | --- | --- | --- | --- | --- |
| Patient 1 | $N_{cell}$ | 877077 | 949845 | 1096565 | 1191979 | 1345744 |
|  | WSS(Pa) | 9.892 | 10.042 | 10.171 | 10.205 | 10.224 |
| Patient 2 | $N_{cell}$ | 590529 | 663147 | 737617 | 812777 | 882729 |
|  | WSS(Pa) | 3.441 | 3.467 | 3.573 | 3.578 | 3.591 |
| Patient 3 | $N_{cell}$ | 883425 | 943711 | 1118507 | 1157100 | 1118507 |
|  | WSS(Pa) | 0.854 | 0.873 | 0.945 | 1.052 | 1.081 |
| Patient 4 | $N_{cell}$ | 428607 | 465817 | 525442 | 583707 | 636012 |
|  | WSS(Pa) | 4.879 | 5.103 | 5.018 | 5.056 | 5.077 |

$N_{cell}:$Number of cell

WSS: Wall shear stress
